# A Genome-Scale Metabolic Model for the Smut-Fungus *Ustilago maydis*

**DOI:** 10.1101/2022.03.03.482780

**Authors:** Ulf W. Liebal, Lena Ullmann, Christian Lieven, Philipp Kohl, Daniel Wibberg, Thiemo Zambanini, Lars M. Blank

## Abstract

*Ustilago maydis* is an important plant pathogen causing corn-smut disease and an effective biotechnological production host. The lack of a comprehensive metabolic overview hinders a full understanding of environmental adaptation and a full use of the organism’s metabolic potential. Here, we report the first genome scale metabolic model (GSMM) of *Ustilago maydis* (iUma22) for the simulation of metabolic activities. iUma22 was reconstructed from sequencing and annotation using PathwayTools, the biomass equation was derived from literature values and from the codon composition. The final model contains over 25% of annotated genes in the sequenced genome. Substrate utilization was corrected by Biolog-Phenotype arrays and exponential batch cultivations were used to test growth predictions. A pan-genome of four different *U. maydis* strains revealed missing metabolic pathways in iUma22. The majority of metabolic differences between iUma22 and the pangenome occurs in the inositol, purine and starch metabolic pathways. The new model allows studies of metabolic adaptations to different environmental niches as well as for biotechnological applications.

## 1. Introduction

*Ustilago maydis* is a model organism and economically important fungus from the division of *Basidiomycota*. The associated corn smut disease affects maize harvest but is also used as food itself [1]. As a parasite, *U. maydis* is growing into the plant tissue to extract substrates for its own metabolic activity. *Ustilaginaceae* show a versatile product spectrum such as organic acids (e.g., itaconate, malate, succinate), polyols (e.g., erythritol, mannitol), and extracellular glycolipids, which are considered value-added chemicals with potential applications in the pharmaceutical, food, and chemical industries. *U. maydis* has developed an effective native production of itaconic acid, an important platform chemical. Indeed, the itaconic acid production in *U. maydis* was improved to surpass the current biotechnological route of *Aspergillus terreus* based production. The advantages are yeast-like growth, high productivities, yields and titer and reduced byproduct formation [2–4], and as model organism, efficient genetic tools are available [5].

The annotated genome sequence of *Ustilago maydis* strain 512 enabled a deeper understanding of the pathogenic mechanisms as well as the metabolic competencies [6]. Annotated genomes can be used to construct genome-scale metabolic models (GSMM), which serve as a knowledge-base of metabolic capacities and allow rational biotechnological engineering [7]. GSMM can be optimized to identify genetic modifications for metabolic engineering that maximize the production of metabolic intermediates. The optimal biotechnological production routes regarding different organisms and metabolic pathways can be computationally evaluated using their respective GSMM [8]. The performance of metabolic microbial and cross-kingdom interactions can be interrogated, to identify exchange metabolites, community stability and metabolic properties that mark the transitions from mutualism to parasitism [9–11].

Here, we present the first high-quality genome-scale metabolic model for *U. maydis* called iUma22. The initial draft model was constructed by the automated PathwayTools workflow with substantial manual adjustments including pathway adjustments derived from Biolog phenotype arrays with 190 carbon substrates. The model quality was assessed using the community evaluation with Memote and the accuracy of rate predictions was tested with glucose growth experiments. The metabolic capacity of the model iUma22 was compared to a pan-genome of different *U. maydis* strains annotated by means of KAAS and enabling reconstruction of KEGG pathways and BRITE hierarchies.

## 2. Materials and Methods

### 2.1. Draft GSMM from Pathway Tools

The genomic DNA sequence of *Ustilago maydis* (Strain 521 FGSC 9021) [6] was obtained from NCBI’s RefSeq project [12]. A corresponding annotation file was then exported from the MIPS Ustilago Maydis Database via the PEDANT Interface [13]. Using the PathoLogic Tool [14], the sequence and annotation files were parsed and in combinationwith MetaCyc reactions database a new Pathway/Genome Database (PGDB) was created. During pathway cleaning reactions from other taxa are pruned, unless there are enzymes matching to all of the reactions. Additional metabolic activity was identified using the ‘Pathway Hole Filler’ function and sequence information of isoenzymes was used to query the proteome of *U. maydis* via pBlast. Protein sequences were queried on PEDANT, MUMDB, MetaCyc or KEGG [15,16] and manually curated while inconclusive polypeptides as well as those that are involved in signaling and other non-metabolic pathways were discarded.

### 2.2 Strains sequenced, pan-genome, KEGG pathway enrichment

To identify metabolic differences within the *U. maydis* strain family, a pan-genome consisting of five *Ustilago maydis* strains was assembled, including strains 198, 482, 485, 512 [17]. The Nanopore Rapid DNA Sequencing kit (SQK-RAD04, Oxford Nanopore Technologies, Oxford, UK) was used for preparation and sequencing was performed on an Oxford Nanopore GridION Mk1 sequencer using a R9.4.1 flow cell. The Nextera XT DNA Sample Preparation Kit (Illumina, San Diego, CA, USA) was used for whole-genome-shotgun PCR-free libraries from 5 μg of gDNA. The library quality was assessed by an Agilent 2000 Bioanalyzer with Agilent High Sensitivity DNA Kit (Agilent Technologies, Santa Clara, CA, USA) for fragment sizes of 500–1000 bp. Paired end sequencing was performed on the Illumina MiSeq platform (2 × 300 bp, v3 chemistry). Adapters and low-quality reads were removed by an in-house software pipeline prior to polishing as recently described [18]. Run control was based on MinKNOW (Oxford Nanopore Technologies) with the 48 h sequencing run protocol. Base calling was performed offline using Bonito, assembly with canu v2.1.1 [19], contigs were polished with Pilon [20] for ten iterative cycles, and for read mapping BWA-MEM [21] and Bowtie2 v2.3.2 [22] in the first and second five iterations, respectively.

Genes were predicted using GeneMark-ES 4.6.2. [23], and functionally annotated using a modified version of the genome annotation platform GenDB 2.0 [24] for eukaryotic genomes [25]. Similarity searches were conducted against COG [26], KEGG [16] and SWISS-PROT [27]. Identification of putative tRNA genes was conducted with tRNAscanSE [28]. Completeness, contamination, and strain heterogeneity were estimated with BUSCO (v3.0.2 [29]), using the fungi-specific single-copy marker genes database (odb9). The obtained genome sequences are compared and documented in more detail in Ullmann et al. (2022) [17]. The pan-genome of all available *U. maydis* strains was calculated by means of EDGAR 3.0 [30]. The KEGG pathway annotation was performed by comparison of E.C. numbers in the pan-genome annotation and E.C. numbers in the reaction description of iUma22. The comparison resulted in three lists, E.C. numbers only present in iUma22 (iUmaNOTpan), present in the pan-genome and iUma (iUmaANDpan), and only present in pan-genome (panNOTiUma). The panNOTiUma list was exported as fasta-file and KAAS [31] was used to annotate the list with KEGG pathway information. The annotation of the genes in the SBML-file was achieved with the BioServices Python package [32].

### 2.3 Biomass equation and growth/non-growth maintenance

The composition of proteins, RNA and DNA was estimated based on the protein and genome sequence respectively, whereas the composition of lipids and the cell wall were results mined from scientific articles [33,34]. The exact biomass composition of *U. maydis* is not available, however, the specific elemental composition [35] and the biomass composition for fungi in general [36] was used as a starting point and linear programming was applied to approximate the total biomass composition (Supplementary 2). The composition values of each monomer were converted into stoichiometric values [37]. For example, to determine the AA composition contribution (in mol_AA_/g_Prot_), first the AA-protein molarity (MP_AA_ in g_AA_/mol_Prot_) was calculated by multiplying the AA codon frequency with the AA molar mass (minus the molar mass of water released during polymerization) (Eq. 1). Normalizing each AA-protein molarity by the overall sum yields the weight fraction of each AA (WP_AA_) (Eq. 2). Division of the AA weight fraction (WP_AA_) by its molar mass and multiplication with the weight fraction of protein to the dry weight (X in g_Prot_/g_CDW_) and conversion from mol to mmol (factor 1000) gives the stoichiometric factor (SF_AA_) (Eq. 3). To calculate the stoichiometric factor of an AA (*SF*_*AA*_) the molar percentage (*MP*) is multiplied with the fractional protein mass per biomass (*X*) (Eq. 2).

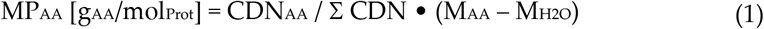

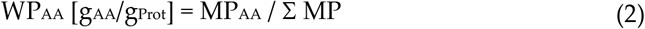

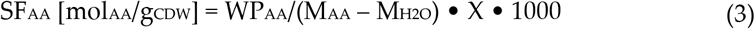

The fraction of protein on total biomass (*X*) is unknown and was determined by linear optimization. The average elemental composition of each macromolecule (protein, DNA., RNA, lipid, cell wall) was determined by summing up the products of the absolute amount of each element. For each macromolecule the C-mole content was calculated by division with the carbon elemental composition [35]. Phosphorous and Sulfur were added from the elemental composition of *S. cerevisiae*. The optimization followed the formula:

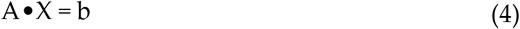

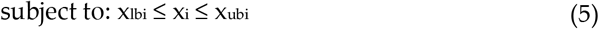

The rows of matrix A correspond to the six individual elements *C, H, O, N, P, S* whereas the columns correspond to the five types of macromolecules (protein, DNA, RNA, lipids, cell wall). The vector b represents the measured elemental biomass *C, H, O, N* supplemented by the elemental content of *P* and *S* of *S. cerevisiae*. The equation was solved for vector *X*, the biomass fractions of each of the macromolecules (Table 1). The lower (x_lb_) and upper (x_ub_) boundary values are provided in Supplementary 2. The ATP-associated maintenance values were set to 46.3 molATP/gCDW and 1.9 molATP /gCDW/h for GAM and NGAM, respectively.

**Table 1.** Macromolecular composition of *U. maydis* calculated by linear optimization. The full composition is provided as Supplementary 1.

| Component | Protein | DNA | RNA | Lipids | Cell Wall |
| --- | --- | --- | --- | --- | --- |
| g/100 gCDW | 30 | 0.3 | 10 | 40 | 16 |

### 2.4 Substrate and growth experiments

For the substrate utilization experiments the Biolog Phenotype Microplates™ PM1 and PM2A were used with *Ustilago maydis* strain 521. Cultures were first grown on YEPS-agar plates at 30°C for at least 24 hours. To prepare the pre-cultures 25 mL of YEPS medium were inoculated from the plates of each strain then performed in 100 mL Erlenmeyer flasks and incubated at 30°C, 200 rpm for 24 hours. (Ecotron Incubation shaker, Infors HT AG, Switzerland). The inoculation fluid was prepared with IFY-0 (1.2x), cell suspension and sterile water to obtain a starting turbidity of 62% T, with 100 µL for each well. The inoculated plates were shaken at 200 rpm with a shaking diameter of 50 mm, at 30°C and with a humidity of 70% up to 168 hours (Multitron Incubation shaker, Infors HT AG, Switzerland). Microbial growth was measured with the SynergyMX (BioTek Instruments, USA) with the optical density at 600 nm. The Biolog raw data is available in Supplementary 4 (PM1) and 5 (PM2A).

The threshold for positive growth was determined by examining the OD histograms for each plate. A normal distribution at low OD values represents the OD range below positive growth (Figure 2). The final growth threshold of 0.4 a.u. was empirically determined to maximize logic consistency and to minimize the integration of false positive metabolic activity. The value approximates the end of a normal distribution of non-growth at low ODs. Experiments conducted for this manuscript and literature data was used to estimate growth rates and glucose uptake rates. The OD measurements of the growth data was converted into g_CDW_/L using the empirical relation from yeast of 0.62 g_CDW_/L /OD (BNID 111182, [38]). The growth rates were identified using a nonlinear fit of the biomass to the Verhulst equation,

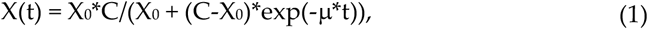

which calculates the biomass from the initial biomass (X_0_), the max biomass capacity (C), the growth rate (m) and time (t). The substrate uptake rate was estimated with a linear equation [39].

## 3. Results and discussion

### 3.1. Description of iUma22

The genome scale model for *Ustilago maydis* was constructed based on the genome sequence and annotation of strain 521 [6]. Table 2 shows the number of represented genes, metabolites and reactions in the new reconstruction and a comparison to the community yeast model ([40], version 8.5.0). Whereas the community yeast model is more comprehensive, iUma22 has a higher gene-to-reaction ratio as well as gene-protein-reaction relationships (GPR), as we aimed to include well connected metabolic pathways with (predicted) annotated genes. *U. maydis* and *S. cerevisiae* have similar number of predicted genes, and when assuming the yeast 8.5.0 as a benchmark of metabolic representation, the iUma22 has reached 70% of genes completeness. There are likely gaps in the secondary metabolism discussed in Ullmann et al. (2022) [17] as well as adaptations to the pathogenic life style.

**Table 2.**
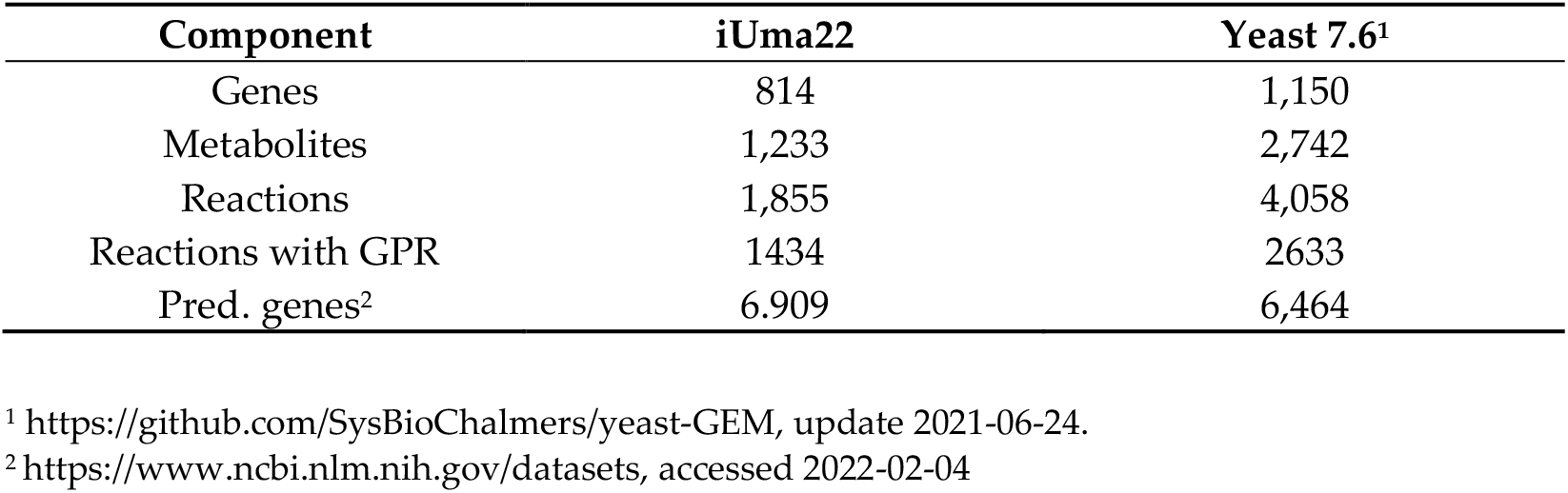
Number of metabolites, reactions and genes of the genome scale metabolic model of *U. maydis* iUma22 and in comparison, the community yeast model (8.5.0 [40]).

| Component | iUma22 | Yeast 7.6<sup>1</sup> |
| --- | --- | --- |
| Genes | 814 | 1,150 |
| Metabolites | 1,233 | 2,742 |
| Reactions | 1,855 | 4,058 |
| Reactions with GPR | 1434 | 2633 |
| Pred. genes<sup>2</sup> | 6.909 | 6,464 |

<sup>1</sup> https://github.com/SysBioChalmers/yeast-GEM, update 2021-06-24.
<sup>2</sup> https://www.ncbi.nlm.nih.gov/datasets, accessed 2022-02-04

The quality of iUma22 was tested with Memote with an overall performance of 42% [41] (Figure 1). Mass and charge balance as well as metabolite connectivity show high quality with scores of over 98%. Memote detects unbounded fluxes that reach boundary conditions during flux variability analysis for 203 reactions on standard media. The stoichiometric consistency of the model could not be evaluated thus decreasing the overall consistency quality to 53%. Note, however, that also for the *S. cerevisiae* community model ([40], version 8.5.0) the stoichiometric consistency test fails. Annotations for metabolites, reactions and genes contain detailed unique annotations. The community yeast model, developed since more than a decade, is evaluated by Memote with a score of 65%.

**Figure 1.**
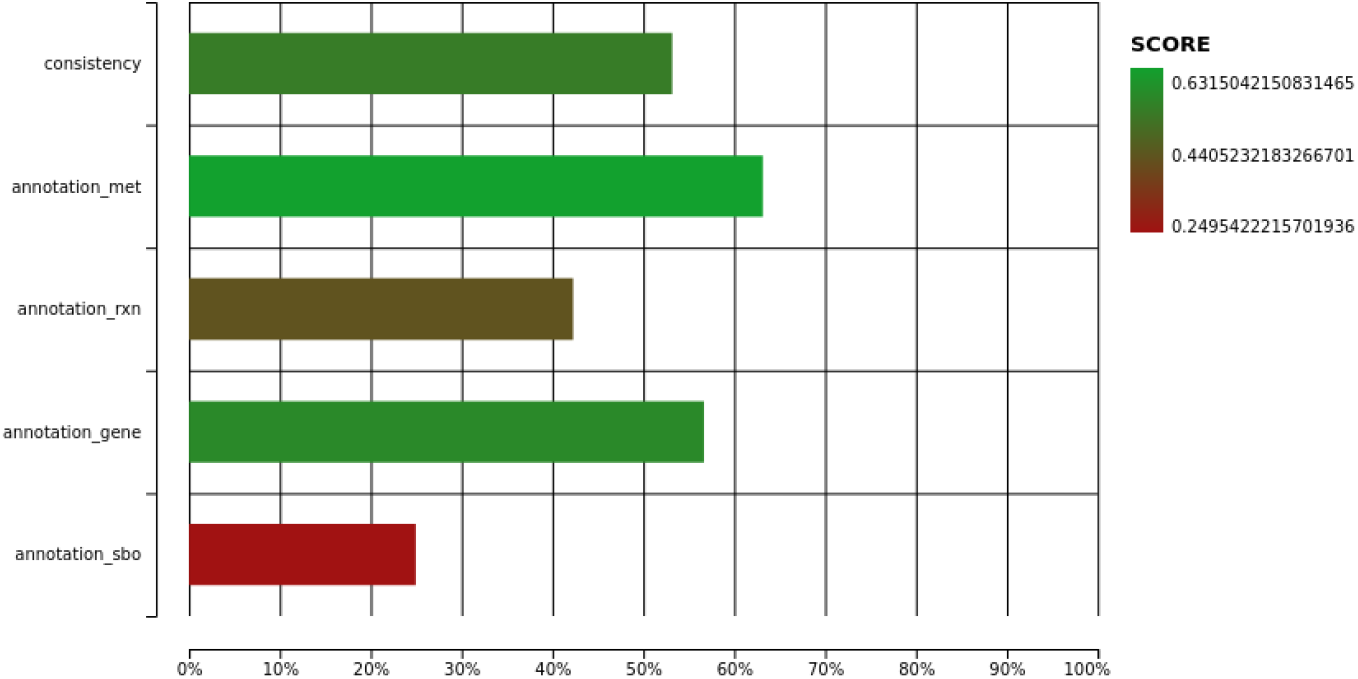
Memote quality report of iUma22 with total score of 42%. The full html report is provided as Supplementary 3.

### 3.2. Carbon Substrate tests with BIOLOG Phenotype arrays

The model iUma22 correctly reproduces 96% of growth phenotypes tested in BIOLOG carbon source assays. We chose an OD threshold of 0.4 a.u. for growth in the plates PM1 and PM2A, a value right after the apparent normal distribution of non-growth for low final OD values (Figure 2). The threshold is a compromise to include growth for glycine dipeptides (PM1: E1, G1, G6, H1), but also includes TCA cycle intermediates (succinate (PM1:A5), fumarate (PM1:F5), aspartate (PM1:A7), malate (PM1:G12)). These TCA cycle intermediates could not be enabled for growth in the model. The largest set of reactions added because of the BIOLOG plates includes di- and oligosaccharide metabolism and methylated central carbon metabolites. Overall, growth took place in 52 wells (34 in PM1 and 19 in PM2A). iUma22 was manually adjusted to reproduce the majority of the growth phenotypes (Figure 2).

**Figure 2.**
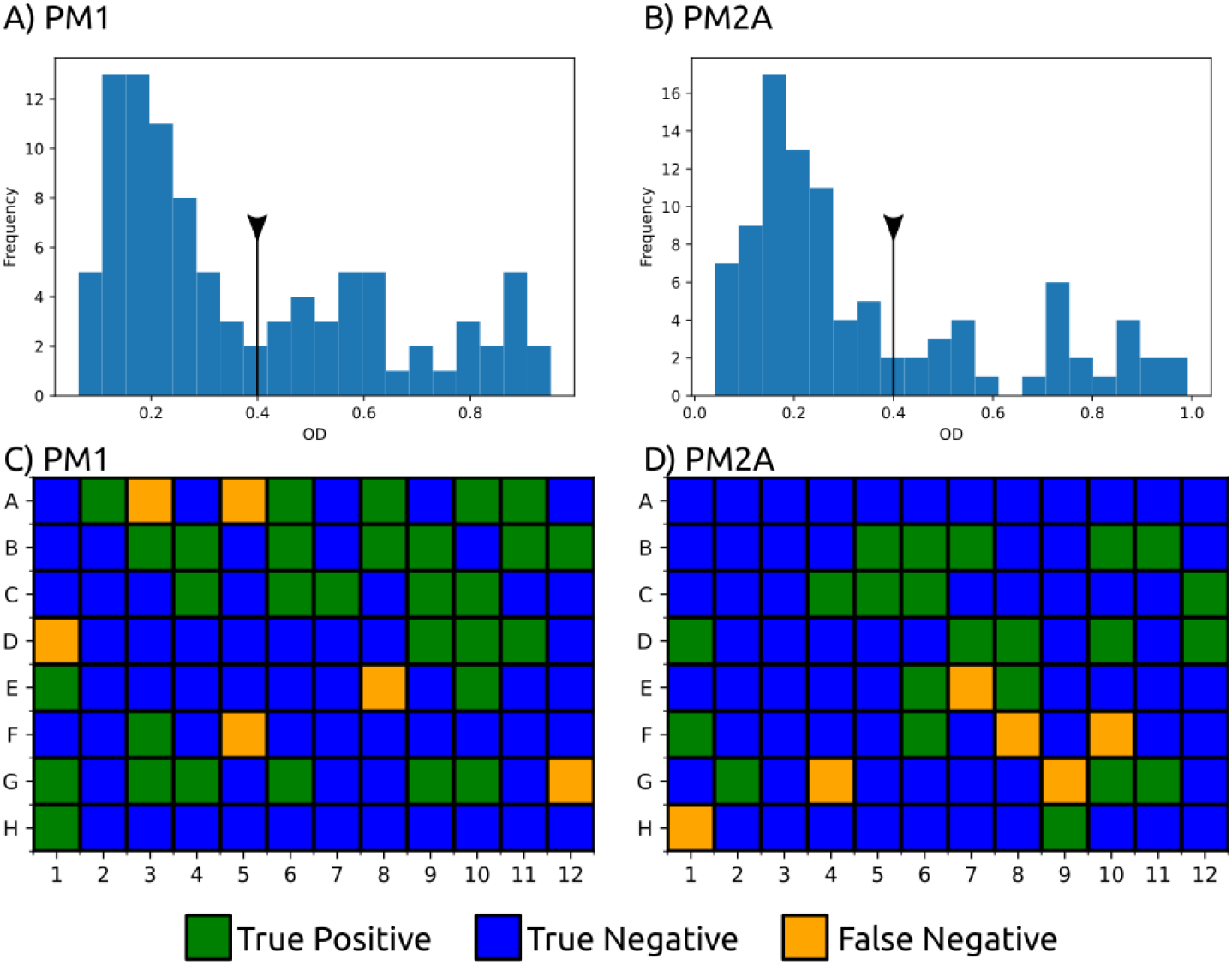
BIOLOG phenotype experiments with carbon sources from PM1 and PM2A. Growth was evaluated by OD 600 after 144h for PM1 (A) and 288h for PM2A (B) with a threshold of 0.4 a.u (black line with triangle) which excludes the normal distribution at low ODs representing no growth. 52 substrates are correctly predicted to growth (true positive, green) and 128 correctly assigned to non-growth by iUma22 (true negative, yellow) in the plates PM1 (C) and PM2A (D). Twelve substrates could not be balanced to enable growth in iUma22 (False Negative). Results of PM1 and PM2A are provided as Supplementary 4 and 5.

While the majority of substrate are correctly reproduced, some metabolites fail to support growth in iUma22. Many intermediates from the TCA cycle do not support growth (2-oxoglutarate, fumarate, succinate, aspartate) and this contrasts for lactate (PM1:B9), malate (PM1:G12) and succinamate (PM2A:F10) in the Biolog results (Figure 2). The degradation of arginine (PM2A:G4), isoleucine (PM2A:G9) and ornithine (H1) is dysfunctional in iUma22, despite the ability of the model to grow on proline, which is a carbon intermediate of the common carbon metabolization following the ornithine-glutamate aminotransferase reaction (E.C. 2.6.1.13, ORNTArm). The metabolization of sebacic acid (PM2A:F8) is not represented in the model, because no information is available in the databases KEGG and Metacyc. Finally, note that false positive results were corrected by removing exchange reactions of the associated metabolites.

### 3.3. Growth rate correlation

We evaluated the growth characteristic of *U. maydis* on glucose and compared the results with predictions of iUma22. We used published batch growth data of *U. maydis* MB215 control strains by Becker et al. (2020) [4], that were genetically modified for optimized biotechnological performance, as well as newly generated data with *U. maydis* strain 521 (Table 3). The growth rate was estimated based on error minimization of the logistic Verhulst growth equation and substrate depletion with a linear equation (Figure 3A and Material and Methods). The yield of each experiment is close to expected and reported yields in *S. cerevisiae* of 0.51 g_CDW_/g_glc_ [42]. Moreover, the correlation between substrate uptake rate and growth rate (Figure 3A) is strong, R2=0.99, and also results in a convincing overall growth associated yield of 0.47 g_CDW_/g_glc_. Note, however, that the highest substrate uptake rate (ID ‘2229v1’, Table 3) is associated with a high standard deviation a higher yield (0.45 g_CDW_/g_glc_), despite the same initial glucose concentration as the experiment by Becker et al. (2020) [4](ID ‘50glc’, Table 3).

**Table 3.**
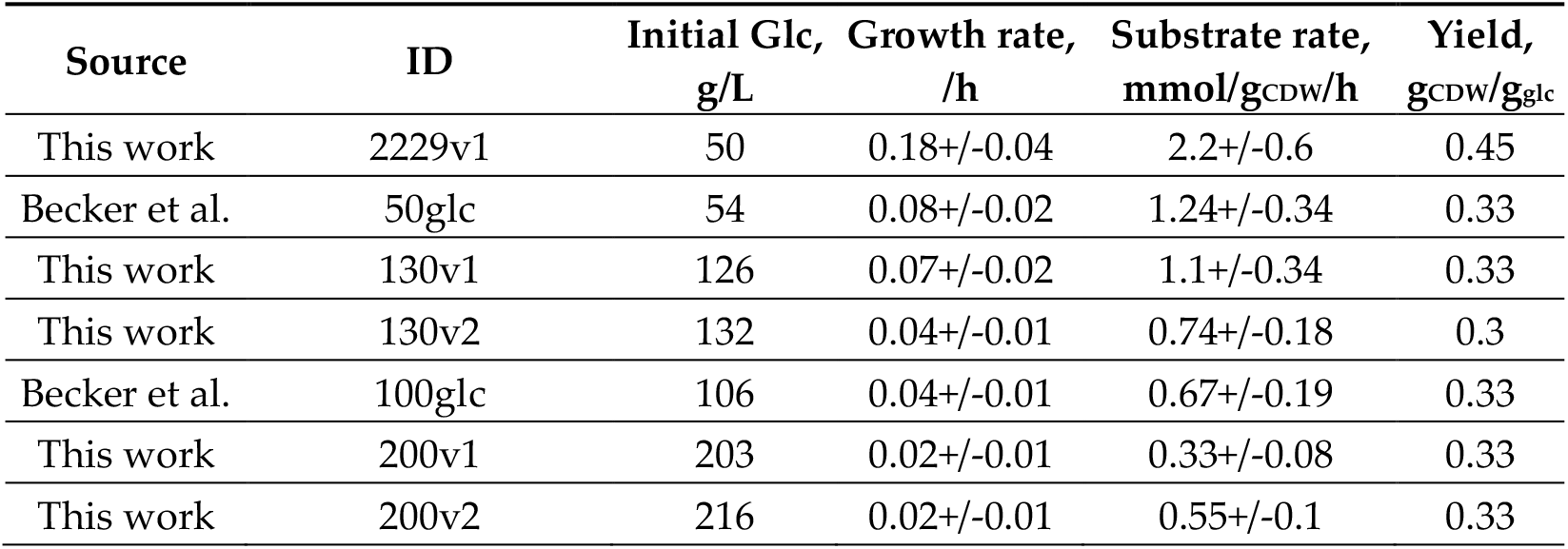
Glucose batch growth experiments were performed and used from the literature [4]. The data provided growth and substrate uptake rates for testing iUma22 predictions. Growth results for each experiment is provided in Supplementary 6.

| Source | ID | Initial Glc,<br>g/L | Growth rate,<br>/h | Substrate rate,<br>mmol/g<sub>CDW</sub>/h | Yield,<br>g<sub>CDW</sub>/g<sub>glc</sub> |
| --- | --- | --- | --- | --- | --- |
| This work | 2229v1 | 50 | 0.18+/-0.04 | 2.2+/-0.6 | 0.45 |
| Becker et al. | 50glc | 54 | 0.08+/-0.02 | 1.24+/-0.34 | 0.33 |
| This work | 130v1 | 126 | 0.07+/-0.02 | 1.1+/-0.34 | 0.33 |
| This work | 130v2 | 132 | 0.04+/-0.01 | 0.74+/-0.18 | 0.3 |
| Becker et al. | 100glc | 106 | 0.04+/-0.01 | 0.67+/-0.19 | 0.33 |
| This work | 200v1 | 203 | 0.02+/-0.01 | 0.33+/-0.08 | 0.33 |
| This work | 200v2 | 216 | 0.02+/-0.01 | 0.55+/-0.1 | 0.33 |

**Figure 3.**
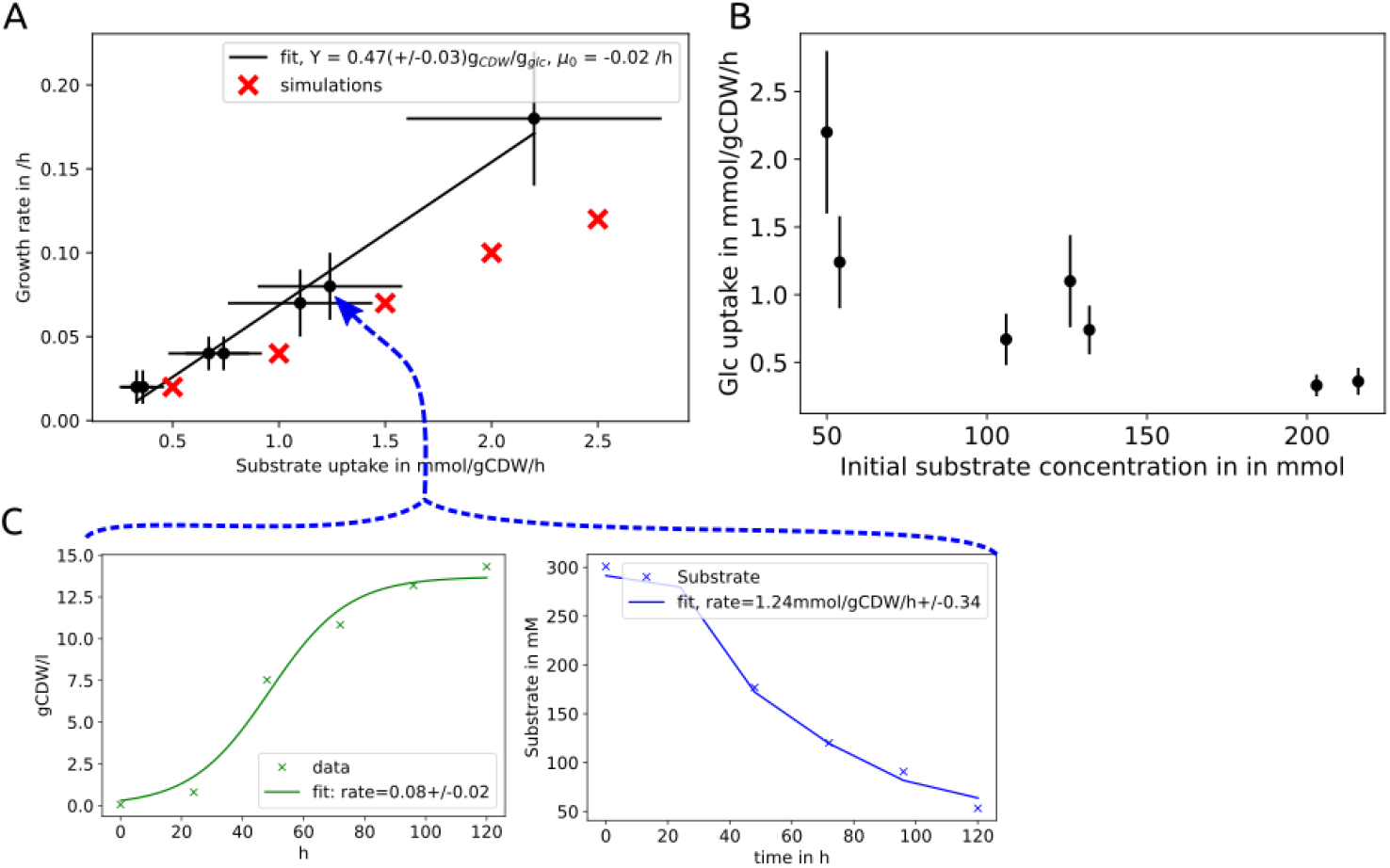
Growth characteristics of *U. maydis* glucose batch cultures and similarity to iUma22 predictions. A) Seven batch experiments on glucose were analyzed to extract growth- and glucose-uptake rates (black dots). B) With increasing initial substrate concentrations the glucose uptake rate decreases. Figures C) and D) represent one example growth experiment of Becker et al. (2020) and the associated analysis. The growth rates were estimated with the Verhulst logarithmic growth model and the growth rate was used to fit a linear equation to identify the substrate uptake rate. The slope of the experimental data in A) provides the biomass yield on glucose with 0.47+/-0.03 g_CDW_/g_glc_, the interception of the x-axis provides the glucose maintenance uptake rate with 0.2+/-0.01 mmol/g_CDW_/h. The growth rate data is provided in Supplementary 6.

The maintenance parameter was calculated as the x-axis interception, i.e., glucose uptake in the absence of growth and results 0.16 mM glc/gcDW/h, similar to *S. cerevisiae* maintenance of 0.2 mM glc/gcDW/h [43]. The growth of *U. maydis* on glucose is substrate inhibited, the higher the glucose concentration the lower is the growth and the substrate uptake rate (Figure 3B, Table 3). Note, that even comparable initial substrate concentrations can result in different substrate uptake and growth dynamics as for the experiments ‘130v1’ with 126 mM and ‘130v2’ with 130 mM which are still consistent according to the comparable yield (Figure 3B, Table 3). The growth predictions by iUma22 coincide to the experiments at low concentrations but underestimate growth at high substrate uptake rates and suggest that the growth associated maintenance parameter should be increased.

### 3.4. U. maydis *Pan-Genome comparison*

We used the available sequenced *U. maydis* strains [17] to compare the enzymatic gene inventory among strains and regarding iUma22. A pan-genome of strains 198, 482, 485 and 512 was constructed by means of EDGAR 3.0 resulting in 7838 coding genes of which 1458 are annotated with an E.C. number. We explored how many genes are shared among all strains and used strain 512 as a reference to identify unique enzyme coding genes and the proportion of shared genes to other strains (Figure 4A). The overwhelming number of genes is shared among all strains (‘all strains’ in Figure 4A), strain 512 has a number of unique enzyme coding genes are more likely shared with other strains because genes with more selective distributions (shared among 3 strains to 1 other strain), are getting less frequent (see also Figure 5 in Ullmann et al., 2022). We then evaluated how the genetic composition of iUma22 differed with respect to the *U. maydis* pan-genome of E.C. annotated genes (Figure 4B). Table 4 shows the top five pathways with the most enzyme annotations for iUma22 unique genes, shared genes and pan-genome unique genes with E.C. numbers. The majority of iUma22 unique genes belong to oxidative phosphorylation, the unique genes in the strain pan-genome belong to diverse central carbon metabolic pathways (Table 4). Particularly noteworthy is the inositol phosphate pathway (Figure 4C), not only because of the highest number of pan-genome unique metabolic capacity but also because inositol was a growth supporting substrate of the Biolog which was manually added to the model.

**Table 4.**
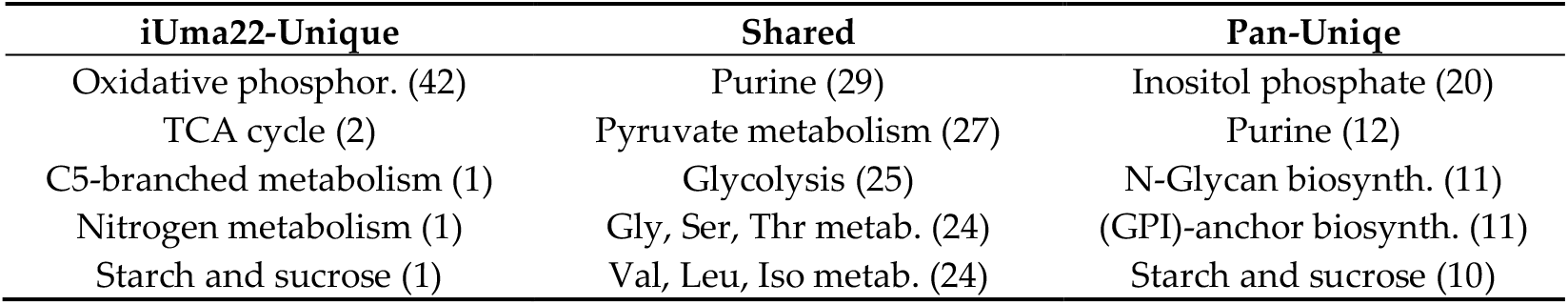
Top five metabolic pathways with the highest number of missing genes in iUma22 compared to the strain Pan genome. The annotation is based on KAAS considering only KEGG pathways (KAAS outputs for iUma22-Unique, Shared and Pan-Unique provided as Supplementary 7-9).

| iUma22-Unique | Shared | Pan-Uniqe |
| --- | --- | --- |
| Oxidative phosphor. (42) | Purine (29) | Inositol phosphate (20) |
| TCA cycle (2) | Pyruvate metabolism (27) | Purine (12) |
| C5-branched metabolism (1) | Glycolysis (25) | N-Glycan biosynth. (11) |
| Nitrogen metabolism (1) | Gly, Ser, Thr metab. (24) | (GPI)-anchor biosynth. (11) |
| Starch and sucrose (1) | Val, Leu, Iso metab. (24) | Starch and sucrose (10) |

**Figure 4.**
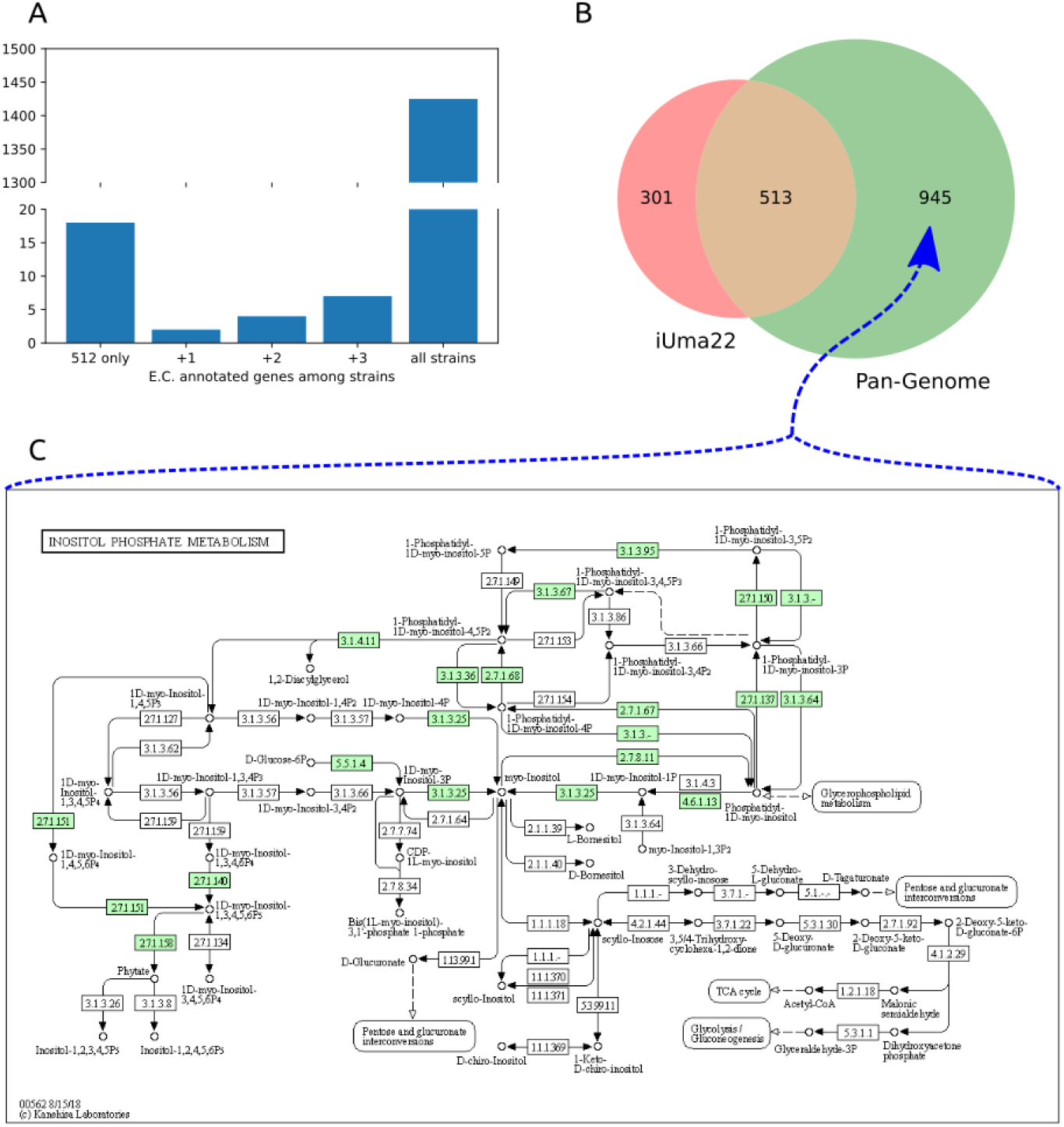
Comparison of enzymes in *U. maydis* strain pan-genome and iUma22. A) E.C. annotated genes in strain 512 that are unique to 512 or shared with the other strains. B) Coverage of the genes in iUma22 of E.C. annotated genes in the pan-genome identified by KAAS. Table 4 show the top five pathways with most association for iUma22 unique, pan-genome unique and their intersection. C) The inositol phosphate metabolism contains with 20 genes the highest level of missing genes in iUma22 (Supplementary 9).

## 5. Conclusions

Here, we present iUma22, a genome scale metabolic model of *U. maydis*, which correctly simulates a large number of substrate phenotypes as well as glucose-based growth rates. The model can be used to identify biotechnological potential of metabolite overproduction and to optimize metabolic engineering strategies. It can also be used to study metabolic shifts in different life cycles of the fungus during plant infection. While the reconstruction was performed based on model strain 521, the genome sequencing of additional *U. maydis* strains provided insight to additional metabolic pathways, which could be used to generate a pan-genome scale metabolic model of *U. maydis*.

## Supplementary Materials

The supplementary material is available on the GitHub page: https://github.com/iAMB-RWTH-Aachen/Ustilago_maydis-GEM

S1: Excel file for calculations of the U. maydis elemental composition.

S2: Matlab workspace file with variables and results of the optimization for the growth equation.

S3: Html output of full SBML quality scan with Memote.

S4: Excel file with the results of Biolog phenotype arrays PM1.

S5: Excel file with the results of Biolog phenotype arrays PM2A.

S6: Excel file with the growth experiments.

S7: Html output of KAAS for pathways unique to iUma22.

S8: Html output of KAAS for pathways shared among iUma22 and strain pan genome.

S9: Html output of KAAS for pathways unique to strain pan genome.

## Author Contributions

Conceptualization, L.M.B., C.L., T.Z., and U.W.L.; methodology, C.L. and U.W.L; experiments L.U. and P.K.; pan-genome construction D.W.; validation, U.W.L. and C.L.; resources, L.M.B.; data curation, U.W.L. and C.L.; writing—original draft preparation, U.W.L.; writing—review and editing, U.W.L., L.U.; visualization, U.W.L.; supervision, T.Z. and U.W.L.; project administration, L.M.B.; funding acquisition, L.M.B. All authors have read and agreed to the published version of the manuscript.

## Funding

This research was funded by the Deutsche Forschungsgemeinschaft (DFG, German Research Foundation) under Germany’s Excellence Strategy within the Cluster of Excellence FSC 2186 “The Fuel Science Center”.

## Data Availability Statement

The data and the model is available on GitHub: https://github.com/iAMB-RWTH-Aachen/Ustilago_maydis-GEM

## Acknowledgments

We appreciate support by Jennifer Scheuplein and Brigida Fabry regarding pathway visualization and Daniel Kaplan on Biolog experiments.

## Conflicts of Interest

The authors declare no conflict of interest. The funders had no role in the design of the study; in the collection, analyses, or interpretation of data; in the writing of the manuscript, or in the decision to publish the results.

## Notes

https://github.com/iAMB-RWTH-Aachen/Ustilago_maydis-GEM

### Competing Interest Statement

The authors have declared no competing interest.

