## Supplemental Data 6 for "A Genome-Scale Metabolic Model for the Smut-Fungus *Ustilago maydis*": Suppl_7_iUma22-Unique.html

KEGG Automatic Annotation Server


### Pathway Mapping (1645025353, query)


- Home
- Help

- [Query list]
- [KO list]
- [BRITE hierarchies]
- [Pathway map]
- [Threshold change]
- [Download KO list]

  

Color Objects in KEGG Pathways *New ver.*

**Default bgcolor:**
 (to highlight the asigned KOs)

**Genes bgcolor:**
 (to change the default color of blue)

00020 Citrate cycle (TCA cycle) (2)

00500 Starch and sucrose metabolism (1)

00660 C5-Branched dibasic acid metabolism (1)

00190 Oxidative phosphorylation (42)

00910 Nitrogen metabolism (1)

00061 Fatty acid biosynthesis (1)

00565 Ether lipid metabolism (1)

03010 Ribosome (2)

04141 Protein processing in endoplasmic reticulum (3)

03050 Proteasome (1)

04010 MAPK signaling pathway (2)

04013 MAPK signaling pathway - fly (1)

04011 MAPK signaling pathway - yeast (3)

04014 Ras signaling pathway (4)

04015 Rap1 signaling pathway (3)

04310 Wnt signaling pathway (2)

04350 TGF-beta signaling pathway (1)

04370 VEGF signaling pathway (2)

04371 Apelin signaling pathway (1)

04072 Phospholipase D signaling pathway (2)

04071 Sphingolipid signaling pathway (2)

04024 cAMP signaling pathway (2)

04022 cGMP-PKG signaling pathway (1)

04151 PI3K-Akt signaling pathway (2)

04150 mTOR signaling pathway (8)

04144 Endocytosis (4)

04145 Phagosome (13)

04142 Lysosome (5)

04140 Autophagy - animal (1)

04138 Autophagy - yeast (1)

04137 Mitophagy - animal (1)

04113 Meiosis - yeast (1)

04510 Focal adhesion (3)

04520 Adherens junction (3)

04530 Tight junction (3)

04810 Regulation of actin cytoskeleton (3)

04611 Platelet activation (1)

04613 Neutrophil extracellular trap formation (2)

04620 Toll-like receptor signaling pathway (1)

04621 NOD-like receptor signaling pathway (1)

04625 C-type lectin receptor signaling pathway (1)

04650 Natural killer cell mediated cytotoxicity (1)

04660 T cell receptor signaling pathway (2)

04662 B cell receptor signaling pathway (1)

04664 Fc epsilon RI signaling pathway (1)

04666 Fc gamma R-mediated phagocytosis (2)

04670 Leukocyte transendothelial migration (3)

04062 Chemokine signaling pathway (4)

04912 GnRH signaling pathway (1)

04921 Oxytocin signaling pathway (1)

04926 Relaxin signaling pathway (1)

04928 Parathyroid hormone synthesis, secretion and action (1)

04260 Cardiac muscle contraction (10)

04270 Vascular smooth muscle contraction (1)

04972 Pancreatic secretion (2)

04978 Mineral absorption (2)

04966 Collecting duct acid secretion (9)

04724 Glutamatergic synapse (1)

04727 GABAergic synapse (2)

04725 Cholinergic synapse (1)

04728 Dopaminergic synapse (1)

04726 Serotonergic synapse (1)

04723 Retrograde endocannabinoid signaling (10)

04721 Synaptic vesicle cycle (12)

04722 Neurotrophin signaling pathway (3)

04744 Phototransduction (1)

04740 Olfactory transduction (1)

04360 Axon guidance (3)

04361 Axon regeneration (2)

04380 Osteoclast differentiation (1)

04713 Circadian entrainment (1)

04714 Thermogenesis (31)

04626 Plant-pathogen interaction (1)

05200 Pathways in cancer (4)

05206 MicroRNAs in cancer (1)

05205 Proteoglycans in cancer (3)

05208 Chemical carcinogenesis - reactive oxygen species (27)

05203 Viral carcinogenesis (4)

05231 Choline metabolism in cancer (1)

05210 Colorectal cancer (2)

05212 Pancreatic cancer (2)

05211 Renal cell carcinoma (2)

05170 Human immunodeficiency virus 1 infection (2)

05171 Coronavirus disease - COVID-19 (2)

05163 Human cytomegalovirus infection (3)

05167 Kaposi sarcoma-associated herpesvirus infection (2)

05169 Epstein-Barr virus infection (2)

05165 Human papillomavirus infection (12)

05110 Vibrio cholerae infection (12)

05120 Epithelial cell signaling in Helicobacter pylori infection (13)

05130 Pathogenic Escherichia coli infection (4)

05132 Salmonella infection (5)

05131 Shigellosis (4)

05135 Yersinia infection (3)

05133 Pertussis (1)

05134 Legionellosis (2)

05152 Tuberculosis (6)

05100 Bacterial invasion of epithelial cells (3)

05146 Amoebiasis (1)

05323 Rheumatoid arthritis (11)

05010 Alzheimer disease (27)

05012 Parkinson disease (27)

05014 Amyotrophic lateral sclerosis (29)

05016 Huntington disease (27)

05017 Spinocerebellar ataxia (1)

05020 Prion disease (28)

05022 Pathways of neurodegeneration - multiple diseases (29)

05032 Morphine addiction (2)

05033 Nicotine addiction (1)

05034 Alcoholism (2)

05417 Lipid and atherosclerosis (3)

05418 Fluid shear stress and atherosclerosis (2)

05415 Diabetic cardiomyopathy (27)

05416 Viral myocarditis (1)

04932 Non-alcoholic fatty liver disease (22)

04933 AGE-RAGE signaling pathway in diabetic complications (2)

01524 Platinum drug resistance (1)

  

- [Query list]
- [KO list]
- [BRITE hierarchies]
- [Pathway map]
- [Threshold change]
- [Download KO list]

- Feedback
- KEGG2
- KEGG
- GenomeNet
- Kanehisa Lab.
