## Supplemental Data 7 for "A Genome-Scale Metabolic Model for the Smut-Fungus *Ustilago maydis*": Suppl_8_Shared.html

KEGG Automatic Annotation Server


### Pathway Mapping (1645026564, query)


- Home
- Help

- [Query list]
- [KO list]
- [BRITE hierarchies]
- [Pathway map]
- [Threshold change]
- [Download KO list]

  

Color Objects in KEGG Pathways *New ver.*

**Default bgcolor:**
 (to highlight the asigned KOs)

**Genes bgcolor:**
 (to change the default color of blue)

00010 Glycolysis / Gluconeogenesis (25)

00020 Citrate cycle (TCA cycle) (19)

00030 Pentose phosphate pathway (18)

00040 Pentose and glucuronate interconversions (7)

00051 Fructose and mannose metabolism (14)

00052 Galactose metabolism (12)

00053 Ascorbate and aldarate metabolism (8)

00500 Starch and sucrose metabolism (13)

00520 Amino sugar and nucleotide sugar metabolism (19)

00620 Pyruvate metabolism (27)

00630 Glyoxylate and dicarboxylate metabolism (17)

00640 Propanoate metabolism (18)

00650 Butanoate metabolism (14)

00660 C5-Branched dibasic acid metabolism (5)

00562 Inositol phosphate metabolism (3)

00190 Oxidative phosphorylation (14)

00710 Carbon fixation in photosynthetic organisms (16)

00720 Carbon fixation pathways in prokaryotes (8)

00680 Methane metabolism (15)

00910 Nitrogen metabolism (8)

00920 Sulfur metabolism (11)

00061 Fatty acid biosynthesis (4)

00062 Fatty acid elongation (5)

00071 Fatty acid degradation (13)

00100 Steroid biosynthesis (12)

00140 Steroid hormone biosynthesis (2)

00561 Glycerolipid metabolism (14)

00564 Glycerophospholipid metabolism (18)

00565 Ether lipid metabolism (3)

00600 Sphingolipid metabolism (5)

00592 alpha-Linolenic acid metabolism (2)

01040 Biosynthesis of unsaturated fatty acids (6)

00230 Purine metabolism (29)

00240 Pyrimidine metabolism (17)

00250 Alanine, aspartate and glutamate metabolism (23)

00260 Glycine, serine and threonine metabolism (24)

00270 Cysteine and methionine metabolism (23)

00280 Valine, leucine and isoleucine degradation (24)

00290 Valine, leucine and isoleucine biosynthesis (10)

00300 Lysine biosynthesis (10)

00310 Lysine degradation (12)

00220 Arginine biosynthesis (14)

00330 Arginine and proline metabolism (14)

00340 Histidine metabolism (8)

00350 Tyrosine metabolism (9)

00360 Phenylalanine metabolism (8)

00380 Tryptophan metabolism (15)

00400 Phenylalanine, tyrosine and tryptophan biosynthesis (14)

00410 beta-Alanine metabolism (11)

00430 Taurine and hypotaurine metabolism (3)

00440 Phosphonate and phosphinate metabolism (2)

00450 Selenocompound metabolism (4)

00460 Cyanoamino acid metabolism (2)

00470 D-Amino acid metabolism (1)

00480 Glutathione metabolism (10)

00510 N-Glycan biosynthesis (9)

00513 Various types of N-glycan biosynthesis (4)

00514 Other types of O-glycan biosynthesis (1)

00604 Glycosphingolipid biosynthesis - ganglio series (1)

00541 O-Antigen nucleotide sugar biosynthesis (5)

00730 Thiamine metabolism (2)

00740 Riboflavin metabolism (5)

00750 Vitamin B6 metabolism (3)

00760 Nicotinate and nicotinamide metabolism (7)

00770 Pantothenate and CoA biosynthesis (13)

00780 Biotin metabolism (3)

00790 Folate biosynthesis (6)

00670 One carbon pool by folate (7)

00830 Retinol metabolism (2)

00860 Porphyrin metabolism (8)

00130 Ubiquinone and other terpenoid-quinone biosynthesis (5)

00900 Terpenoid backbone biosynthesis (8)

00909 Sesquiterpenoid and triterpenoid biosynthesis (2)

00906 Carotenoid biosynthesis (1)

00981 Insect hormone biosynthesis (1)

00903 Limonene and pinene degradation (1)

00281 Geraniol degradation (2)

01051 Biosynthesis of ansamycins (1)

00940 Phenylpropanoid biosynthesis (2)

00950 Isoquinoline alkaloid biosynthesis (2)

00960 Tropane, piperidine and pyridine alkaloid biosynthesis (3)

00232 Caffeine metabolism (2)

00966 Glucosinolate biosynthesis (1)

00311 Penicillin and cephalosporin biosynthesis (1)

00332 Carbapenem biosynthesis (2)

00261 Monobactam biosynthesis (3)

00521 Streptomycin biosynthesis (2)

00524 Neomycin, kanamycin and gentamicin biosynthesis (1)

00401 Novobiocin biosynthesis (1)

00405 Phenazine biosynthesis (1)

00333 Prodigiosin biosynthesis (1)

00254 Aflatoxin biosynthesis (1)

00999 Biosynthesis of various plant secondary metabolites (3)

00362 Benzoate degradation (5)

00627 Aminobenzoate degradation (4)

00625 Chloroalkane and chloroalkene degradation (3)

00643 Styrene degradation (3)

00791 Atrazine degradation (1)

00930 Caprolactam degradation (4)

00621 Dioxin degradation (1)

00626 Naphthalene degradation (3)

00624 Polycyclic aromatic hydrocarbon degradation (1)

00980 Metabolism of xenobiotics by cytochrome P450 (2)

00982 Drug metabolism - cytochrome P450 (2)

00983 Drug metabolism - other enzymes (10)

03022 Basal transcription factors (1)

03040 Spliceosome (1)

00970 Aminoacyl-tRNA biosynthesis (5)

03013 Nucleocytoplasmic transport (1)

03015 mRNA surveillance pathway (1)

04141 Protein processing in endoplasmic reticulum (1)

04122 Sulfur relay system (2)

03018 RNA degradation (3)

03410 Base excision repair (1)

03420 Nucleotide excision repair (1)

02020 Two-component system (4)

04013 MAPK signaling pathway - fly (1)

04016 MAPK signaling pathway - plant (2)

04011 MAPK signaling pathway - yeast (2)

04014 Ras signaling pathway (1)

04066 HIF-1 signaling pathway (7)

04068 FoxO signaling pathway (1)

04020 Calcium signaling pathway (1)

04070 Phosphatidylinositol signaling system (1)

04072 Phospholipase D signaling pathway (2)

04071 Sphingolipid signaling pathway (5)

04024 cAMP signaling pathway (4)

04022 cGMP-PKG signaling pathway (2)

04152 AMPK signaling pathway (6)

04150 mTOR signaling pathway (1)

04144 Endocytosis (1)

04145 Phagosome (1)

04142 Lysosome (2)

04146 Peroxisome (12)

04138 Autophagy - yeast (2)

04111 Cell cycle - yeast (1)

04216 Ferroptosis (3)

04217 Necroptosis (2)

04115 p53 signaling pathway (1)

02024 Quorum sensing (5)

02025 Biofilm formation - Pseudomonas aeruginosa (1)

04622 RIG-I-like receptor signaling pathway (1)

04666 Fc gamma R-mediated phagocytosis (1)

04911 Insulin secretion (1)

04910 Insulin signaling pathway (3)

04922 Glucagon signaling pathway (7)

04923 Regulation of lipolysis in adipocytes (1)

04920 Adipocytokine signaling pathway (1)

03320 PPAR signaling pathway (8)

04912 GnRH signaling pathway (1)

04917 Prolactin signaling pathway (1)

04918 Thyroid hormone synthesis (2)

04919 Thyroid hormone signaling pathway (3)

04928 Parathyroid hormone synthesis, secretion and action (1)

04925 Aldosterone synthesis and secretion (1)

04260 Cardiac muscle contraction (3)

04261 Adrenergic signaling in cardiomyocytes (2)

04970 Salivary secretion (1)

04971 Gastric acid secretion (1)

04972 Pancreatic secretion (2)

04976 Bile secretion (2)

04973 Carbohydrate digestion and absorption (2)

04974 Protein digestion and absorption (1)

04975 Fat digestion and absorption (4)

04979 Cholesterol metabolism (1)

04978 Mineral absorption (2)

04960 Aldosterone-regulated sodium reabsorption (1)

04961 Endocrine and other factor-regulated calcium reabsorption (1)

04964 Proximal tubule bicarbonate reclamation (1)

04966 Collecting duct acid secretion (1)

04724 Glutamatergic synapse (2)

04727 GABAergic synapse (3)

04723 Retrograde endocannabinoid signaling (8)

04721 Synaptic vesicle cycle (1)

04211 Longevity regulating pathway (1)

04212 Longevity regulating pathway - worm (3)

04213 Longevity regulating pathway - multiple species (1)

04714 Thermogenesis (14)

04626 Plant-pathogen interaction (1)

05200 Pathways in cancer (2)

05202 Transcriptional misregulation in cancer (1)

05205 Proteoglycans in cancer (1)

05208 Chemical carcinogenesis - reactive oxygen species (14)

05203 Viral carcinogenesis (1)

05230 Central carbon metabolism in cancer (8)

05231 Choline metabolism in cancer (2)

05212 Pancreatic cancer (1)

05211 Renal cell carcinoma (1)

05166 Human T-cell leukemia virus 1 infection (1)

05164 Influenza A (1)

05165 Human papillomavirus infection (2)

05110 Vibrio cholerae infection (1)

05120 Epithelial cell signaling in Helicobacter pylori infection (1)

05130 Pathogenic Escherichia coli infection (1)

05132 Salmonella infection (1)

05131 Shigellosis (1)

05146 Amoebiasis (1)

05143 African trypanosomiasis (1)

05323 Rheumatoid arthritis (1)

05010 Alzheimer disease (13)

05012 Parkinson disease (11)

05014 Amyotrophic lateral sclerosis (11)

05016 Huntington disease (12)

05017 Spinocerebellar ataxia (1)

05020 Prion disease (11)

05022 Pathways of neurodegeneration - multiple diseases (12)

05417 Lipid and atherosclerosis (1)

05418 Fluid shear stress and atherosclerosis (1)

05410 Hypertrophic cardiomyopathy (1)

05412 Arrhythmogenic right ventricular cardiomyopathy (1)

05414 Dilated cardiomyopathy (1)

05415 Diabetic cardiomyopathy (18)

04930 Type II diabetes mellitus (2)

04940 Type I diabetes mellitus (1)

04936 Alcoholic liver disease (7)

04932 Non-alcoholic fatty liver disease (10)

04931 Insulin resistance (1)

04934 Cushing syndrome (1)

01524 Platinum drug resistance (1)

01523 Antifolate resistance (5)

  

- [Query list]
- [KO list]
- [BRITE hierarchies]
- [Pathway map]
- [Threshold change]
- [Download KO list]

- Feedback
- KEGG2
- KEGG
- GenomeNet
- Kanehisa Lab.
