## Supplemental Data 8 for "A Genome-Scale Metabolic Model for the Smut-Fungus *Ustilago maydis*": Suppl_9_Pan-Unique.html

KEGG Automatic Annotation Server


### Pathway Mapping (1644568906, query)


- Home
- Help

- [Query list]
- [KO list]
- [BRITE hierarchies]
- [Pathway map]
- [Threshold change]
- [Download KO list]

  

Color Objects in KEGG Pathways *New ver.*

**Default bgcolor:**
 (to highlight the asigned KOs)

**Genes bgcolor:**
 (to change the default color of blue)

00010 Glycolysis / Gluconeogenesis (3)

00020 Citrate cycle (TCA cycle) (2)

00030 Pentose phosphate pathway (1)

00040 Pentose and glucuronate interconversions (5)

00051 Fructose and mannose metabolism (4)

00052 Galactose metabolism (4)

00053 Ascorbate and aldarate metabolism (2)

00500 Starch and sucrose metabolism (10)

00520 Amino sugar and nucleotide sugar metabolism (7)

00620 Pyruvate metabolism (3)

00630 Glyoxylate and dicarboxylate metabolism (4)

00640 Propanoate metabolism (2)

00650 Butanoate metabolism (2)

00562 Inositol phosphate metabolism (20)

00190 Oxidative phosphorylation (1)

00910 Nitrogen metabolism (1)

00920 Sulfur metabolism (3)

00062 Fatty acid elongation (2)

00071 Fatty acid degradation (3)

00073 Cutin, suberine and wax biosynthesis (2)

00100 Steroid biosynthesis (3)

00120 Primary bile acid biosynthesis (2)

00561 Glycerolipid metabolism (7)

00564 Glycerophospholipid metabolism (8)

00565 Ether lipid metabolism (1)

00600 Sphingolipid metabolism (9)

00590 Arachidonic acid metabolism (4)

01040 Biosynthesis of unsaturated fatty acids (2)

00230 Purine metabolism (12)

00240 Pyrimidine metabolism (6)

00250 Alanine, aspartate and glutamate metabolism (3)

00260 Glycine, serine and threonine metabolism (5)

00270 Cysteine and methionine metabolism (5)

00280 Valine, leucine and isoleucine degradation (2)

00310 Lysine degradation (6)

00220 Arginine biosynthesis (1)

00330 Arginine and proline metabolism (8)

00340 Histidine metabolism (4)

00350 Tyrosine metabolism (4)

00360 Phenylalanine metabolism (5)

00380 Tryptophan metabolism (5)

00400 Phenylalanine, tyrosine and tryptophan biosynthesis (1)

00410 beta-Alanine metabolism (3)

00430 Taurine and hypotaurine metabolism (3)

00450 Selenocompound metabolism (2)

00460 Cyanoamino acid metabolism (4)

00480 Glutathione metabolism (9)

00510 N-Glycan biosynthesis (11)

00513 Various types of N-glycan biosynthesis (8)

00515 Mannose type O-glycan biosynthesis (1)

00514 Other types of O-glycan biosynthesis (2)

00531 Glycosaminoglycan degradation (1)

00563 Glycosylphosphatidylinositol (GPI)-anchor biosynthesis (11)

00603 Glycosphingolipid biosynthesis - globo and isoglobo series (2)

00604 Glycosphingolipid biosynthesis - ganglio series (1)

00541 O-Antigen nucleotide sugar biosynthesis (1)

00511 Other glycan degradation (3)

00730 Thiamine metabolism (6)

00740 Riboflavin metabolism (5)

00750 Vitamin B6 metabolism (4)

00760 Nicotinate and nicotinamide metabolism (9)

00770 Pantothenate and CoA biosynthesis (3)

00780 Biotin metabolism (2)

00785 Lipoic acid metabolism (2)

00790 Folate biosynthesis (6)

00670 One carbon pool by folate (4)

00830 Retinol metabolism (1)

00860 Porphyrin metabolism (6)

00130 Ubiquinone and other terpenoid-quinone biosynthesis (3)

00900 Terpenoid backbone biosynthesis (8)

00906 Carotenoid biosynthesis (1)

00981 Insect hormone biosynthesis (1)

00908 Zeatin biosynthesis (1)

00903 Limonene and pinene degradation (1)

00940 Phenylpropanoid biosynthesis (2)

00901 Indole alkaloid biosynthesis (1)

00950 Isoquinoline alkaloid biosynthesis (3)

00960 Tropane, piperidine and pyridine alkaloid biosynthesis (1)

00965 Betalain biosynthesis (2)

00521 Streptomycin biosynthesis (2)

00999 Biosynthesis of various plant secondary metabolites (1)

00362 Benzoate degradation (3)

00627 Aminobenzoate degradation (3)

00364 Fluorobenzoate degradation (1)

00625 Chloroalkane and chloroalkene degradation (2)

00361 Chlorocyclohexane and chlorobenzene degradation (1)

00623 Toluene degradation (2)

00643 Styrene degradation (2)

00980 Metabolism of xenobiotics by cytochrome P450 (3)

00982 Drug metabolism - cytochrome P450 (3)

00983 Drug metabolism - other enzymes (4)

03020 RNA polymerase (4)

03022 Basal transcription factors (4)

03040 Spliceosome (12)

00970 Aminoacyl-tRNA biosynthesis (22)

03013 Nucleocytoplasmic transport (3)

03015 mRNA surveillance pathway (10)

03008 Ribosome biogenesis in eukaryotes (20)

03060 Protein export (6)

04141 Protein processing in endoplasmic reticulum (24)

04120 Ubiquitin mediated proteolysis (27)

04122 Sulfur relay system (3)

03050 Proteasome (14)

03018 RNA degradation (12)

03030 DNA replication (17)

03410 Base excision repair (15)

03420 Nucleotide excision repair (10)

03430 Mismatch repair (4)

03440 Homologous recombination (9)

03450 Non-homologous end-joining (3)

03460 Fanconi anemia pathway (13)

02010 ABC transporters (2)

03070 Bacterial secretion system (1)

02020 Two-component system (5)

04010 MAPK signaling pathway (8)

04013 MAPK signaling pathway - fly (5)

04016 MAPK signaling pathway - plant (2)

04011 MAPK signaling pathway - yeast (22)

04012 ErbB signaling pathway (5)

04014 Ras signaling pathway (5)

04015 Rap1 signaling pathway (3)

04310 Wnt signaling pathway (7)

04330 Notch signaling pathway (2)

04340 Hedgehog signaling pathway (2)

04341 Hedgehog signaling pathway - fly (3)

04350 TGF-beta signaling pathway (3)

04390 Hippo signaling pathway (4)

04391 Hippo signaling pathway - fly (3)

04392 Hippo signaling pathway - multiple species (4)

04370 VEGF signaling pathway (5)

04371 Apelin signaling pathway (6)

04630 JAK-STAT signaling pathway (2)

04064 NF-kappa B signaling pathway (2)

04668 TNF signaling pathway (3)

04066 HIF-1 signaling pathway (4)

04068 FoxO signaling pathway (9)

04020 Calcium signaling pathway (6)

04070 Phosphatidylinositol signaling system (20)

04072 Phospholipase D signaling pathway (5)

04071 Sphingolipid signaling pathway (11)

04024 cAMP signaling pathway (5)

04022 cGMP-PKG signaling pathway (4)

04151 PI3K-Akt signaling pathway (9)

04152 AMPK signaling pathway (5)

04150 mTOR signaling pathway (8)

04144 Endocytosis (3)

04145 Phagosome (2)

04142 Lysosome (7)

04146 Peroxisome (10)

04140 Autophagy - animal (14)

04138 Autophagy - yeast (19)

04136 Autophagy - other (6)

04137 Mitophagy - animal (4)

04139 Mitophagy - yeast (13)

04110 Cell cycle (18)

04111 Cell cycle - yeast (22)

04112 Cell cycle - Caulobacter (1)

04113 Meiosis - yeast (21)

04114 Oocyte meiosis (9)

04210 Apoptosis (5)

04214 Apoptosis - fly (7)

04217 Necroptosis (2)

04115 p53 signaling pathway (4)

04218 Cellular senescence (13)

04510 Focal adhesion (8)

04520 Adherens junction (3)

04530 Tight junction (3)

04540 Gap junction (3)

04550 Signaling pathways regulating pluripotency of stem cells (4)

02024 Quorum sensing (2)

02025 Biofilm formation - Pseudomonas aeruginosa (1)

04810 Regulation of actin cytoskeleton (5)

04611 Platelet activation (4)

04613 Neutrophil extracellular trap formation (8)

04620 Toll-like receptor signaling pathway (2)

04624 Toll and Imd signaling pathway (3)

04621 NOD-like receptor signaling pathway (4)

04622 RIG-I-like receptor signaling pathway (3)

04623 Cytosolic DNA-sensing pathway (2)

04625 C-type lectin receptor signaling pathway (4)

04650 Natural killer cell mediated cytotoxicity (4)

04612 Antigen processing and presentation (1)

04660 T cell receptor signaling pathway (6)

04658 Th1 and Th2 cell differentiation (3)

04659 Th17 cell differentiation (4)

04657 IL-17 signaling pathway (4)

04662 B cell receptor signaling pathway (3)

04664 Fc epsilon RI signaling pathway (4)

04666 Fc gamma R-mediated phagocytosis (5)

04670 Leukocyte transendothelial migration (2)

04062 Chemokine signaling pathway (4)

04911 Insulin secretion (3)

04910 Insulin signaling pathway (7)

04922 Glucagon signaling pathway (5)

04923 Regulation of lipolysis in adipocytes (2)

04920 Adipocytokine signaling pathway (2)

03320 PPAR signaling pathway (3)

04929 GnRH secretion (2)

04912 GnRH signaling pathway (4)

04913 Ovarian steroidogenesis (1)

04915 Estrogen signaling pathway (3)

04914 Progesterone-mediated oocyte maturation (4)

04917 Prolactin signaling pathway (3)

04921 Oxytocin signaling pathway (7)

04926 Relaxin signaling pathway (4)

04935 Growth hormone synthesis, secretion and action (6)

04918 Thyroid hormone synthesis (4)

04919 Thyroid hormone signaling pathway (10)

04928 Parathyroid hormone synthesis, secretion and action (3)

04916 Melanogenesis (5)

04924 Renin secretion (2)

04614 Renin-angiotensin system (2)

04925 Aldosterone synthesis and secretion (4)

04927 Cortisol synthesis and secretion (1)

04260 Cardiac muscle contraction (1)

04261 Adrenergic signaling in cardiomyocytes (7)

04270 Vascular smooth muscle contraction (4)

04970 Salivary secretion (4)

04971 Gastric acid secretion (3)

04972 Pancreatic secretion (3)

04976 Bile secretion (3)

04973 Carbohydrate digestion and absorption (2)

04974 Protein digestion and absorption (1)

04979 Cholesterol metabolism (2)

04977 Vitamin digestion and absorption (1)

04978 Mineral absorption (2)

04962 Vasopressin-regulated water reabsorption (2)

04960 Aldosterone-regulated sodium reabsorption (4)

04961 Endocrine and other factor-regulated calcium reabsorption (3)

04964 Proximal tubule bicarbonate reclamation (1)

04724 Glutamatergic synapse (4)

04727 GABAergic synapse (5)

04725 Cholinergic synapse (3)

04728 Dopaminergic synapse (8)

04726 Serotonergic synapse (4)

04720 Long-term potentiation (5)

04730 Long-term depression (3)

04723 Retrograde endocannabinoid signaling (6)

04721 Synaptic vesicle cycle (1)

04722 Neurotrophin signaling pathway (4)

04745 Phototransduction - fly (1)

04740 Olfactory transduction (1)

04742 Taste transduction (1)

04750 Inflammatory mediator regulation of TRP channels (4)

04320 Dorso-ventral axis formation (1)

04360 Axon guidance (6)

04361 Axon regeneration (6)

04380 Osteoclast differentiation (3)

04211 Longevity regulating pathway (4)

04212 Longevity regulating pathway - worm (8)

04213 Longevity regulating pathway - multiple species (8)

04710 Circadian rhythm (1)

04713 Circadian entrainment (3)

04711 Circadian rhythm - fly (1)

04712 Circadian rhythm - plant (1)

04714 Thermogenesis (4)

05200 Pathways in cancer (11)

05202 Transcriptional misregulation in cancer (4)

05206 MicroRNAs in cancer (8)

05205 Proteoglycans in cancer (8)

05204 Chemical carcinogenesis - DNA adducts (3)

05207 Chemical carcinogenesis - receptor activation (6)

05208 Chemical carcinogenesis - reactive oxygen species (9)

05203 Viral carcinogenesis (10)

05230 Central carbon metabolism in cancer (3)

05231 Choline metabolism in cancer (5)

05235 PD-L1 expression and PD-1 checkpoint pathway in cancer (6)

05210 Colorectal cancer (4)

05212 Pancreatic cancer (3)

05225 Hepatocellular carcinoma (8)

05226 Gastric cancer (6)

05214 Glioma (6)

05216 Thyroid cancer (2)

05221 Acute myeloid leukemia (2)

05220 Chronic myeloid leukemia (3)

05217 Basal cell carcinoma (2)

05218 Melanoma (3)

05211 Renal cell carcinoma (3)

05219 Bladder cancer (1)

05215 Prostate cancer (5)

05213 Endometrial cancer (5)

05224 Breast cancer (5)

05222 Small cell lung cancer (2)

05223 Non-small cell lung cancer (4)

05166 Human T-cell leukemia virus 1 infection (12)

05170 Human immunodeficiency virus 1 infection (10)

05161 Hepatitis B (4)

05160 Hepatitis C (5)

05171 Coronavirus disease - COVID-19 (3)

05164 Influenza A (2)

05162 Measles (3)

05168 Herpes simplex virus 1 infection (3)

05163 Human cytomegalovirus infection (7)

05167 Kaposi sarcoma-associated herpesvirus infection (6)

05169 Epstein-Barr virus infection (5)

05165 Human papillomavirus infection (10)

05110 Vibrio cholerae infection (2)

05120 Epithelial cell signaling in Helicobacter pylori infection (2)

05130 Pathogenic Escherichia coli infection (3)

05132 Salmonella infection (4)

05131 Shigellosis (12)

05135 Yersinia infection (4)

05133 Pertussis (2)

05134 Legionellosis (1)

05152 Tuberculosis (6)

05146 Amoebiasis (2)

05145 Toxoplasmosis (3)

05140 Leishmaniasis (2)

05142 Chagas disease (3)

05143 African trypanosomiasis (1)

05340 Primary immunodeficiency (1)

05010 Alzheimer disease (25)

05012 Parkinson disease (24)

05014 Amyotrophic lateral sclerosis (25)

05016 Huntington disease (23)

05017 Spinocerebellar ataxia (24)

05020 Prion disease (21)

05022 Pathways of neurodegeneration - multiple diseases (36)

05030 Cocaine addiction (2)

05031 Amphetamine addiction (6)

05032 Morphine addiction (2)

05034 Alcoholism (9)

05417 Lipid and atherosclerosis (9)

05418 Fluid shear stress and atherosclerosis (2)

05414 Dilated cardiomyopathy (1)

05415 Diabetic cardiomyopathy (9)

04930 Type II diabetes mellitus (2)

04940 Type I diabetes mellitus (1)

04936 Alcoholic liver disease (6)

04932 Non-alcoholic fatty liver disease (3)

04931 Insulin resistance (6)

04933 AGE-RAGE signaling pathway in diabetic complications (4)

04934 Cushing syndrome (4)

01521 EGFR tyrosine kinase inhibitor resistance (5)

01524 Platinum drug resistance (8)

01523 Antifolate resistance (1)

01522 Endocrine resistance (5)

  

- [Query list]
- [KO list]
- [BRITE hierarchies]
- [Pathway map]
- [Threshold change]
- [Download KO list]

- Feedback
- KEGG2
- KEGG
- GenomeNet
- Kanehisa Lab.
